# JNK activity modulates postsynaptic scaffold protein SAP102 and kainate receptor dynamics in dendritic spines

**DOI:** 10.1101/2021.04.30.442109

**Authors:** Stella-Amrei Kunde, Bettina Schmerl, Elham Ahmadyar, Nils Rademacher, Hanna L. Zieger, Sarah A. Shoichet

## Abstract

We show here that the dynamics of the synaptic scaffold molecule SAP102 are negatively regulated by JNK inhibition, that SAP102 is a direct phosphorylation target of JNK3, and that SAP102 regulation by JNK is restricted to neurons that harbour mature synapses. We further demonstrate that SAP102 and JNK3 cooperate in the regulated trafficking of kainate receptors to the cell membrane. Specifically, we observe that SAP102, JNK3, and the kainate receptor subunit GluK2 exhibit overlapping expression at synaptic sites, and that modulating JNK activity influences the surface expression of the kainate receptor subunit GluK2 in a neuronal context. We also show that SAP102 participates in this process in a JNK-dependent fashion. In summary, our data support a model in which JNK-mediated regulation of SAP102 influences the dynamic trafficking of kainate receptors to postsynaptic sites, and thus shed light on common pathophysiological mechanisms underlying the cognitive developmental defects associated with diverse mutations.

## Introduction

Proper network formation relies on the regulated expression of diverse neuronal proteins and also on the coordinated generation and modulation of synaptic connections during development. In excitatory neurons, synaptic function is largely dependent on the integrity of the postsynaptic density (PSD); the molecular composition of the PSD has therefore been studied in considerable depth (for reviews, see *e.g*. ^1^ and ^2^). In addition to neurotransmitter receptors and trans-synaptic cell adhesion molecules at the postsynaptic membrane, scaffold proteins play an integral role in defining the functional architecture of the PSD, and their coordinated expression and regulation is critical for proper synapse formation and maintenance. It follows from this that mutations in diverse postsynaptic scaffold proteins have been implicated in neurodevelopmental disorders (see *e.g*. ^3-5^ for reviews).

Among the postsynaptic scaffold proteins associated with neurodevelopmental disorders, the PSD-95 family of synaptic membrane-associated guanylate kinases (MAGUKs) is of particular importance. PSD-95 family MAGUKS are among the most abundant scaffold proteins at the postsynaptic sites of glutamatergic neurons, and they are recognised as central building blocks of the PSD ^6^. They are critical for coordinating the trafficking and anchoring of glutamate receptors at the postsynaptic membrane^7^, and these functions of MAGUKs are especially important during development^8^. Interestingly, relatively few disease-causing mutations have been found in PSD-95, the prototypical synaptic MAGUK that has been investigated most extensively. Among synaptic MAGUKs, SAP102 is recognised for its high expression during early development and critical role in synapse development and maturation^9,10^. In line with an important developmental function of this particular synaptic MAGUK, numerous monogenic forms of developmental delay have been linked to genetic alterations of SAP102^11-15^ (gene *DLG3*, see also OMIM #300850 MRX90). SAP102 also differs functionally from other synaptic MAGUKs in that it behaves differently in dendritic spines: while PSD-95 is anchored stably at the PSD, SAP102 exhibits a comparatively high mobility into and out of the spines^16^, suggesting a unique role for this MAGUK in receptor trafficking to and from synaptic sites. Moreover, it has been shown that the rate with which SAP102 enters and exits the spine can be influenced by post-translational modifications^17,18^, highlighting that SAP102 is potentially able to modulate its cellular function in response to extracellular signals, which are especially relevant during synapse formation, learning, and excitotoxic stress.

We have previously shown that SAP102 is able to bind to the c-Jun N-terminal kinase JNK3^19^, which is the CNS-specific member of the JNK family of MAP kinases, *i. e*. the terminal kinases in one branch of the MAP kinase cascade that coordinates cellular responses to diverse external stimuli. Like SAP102, JNK3 is also interesting in the context of disease: JNK3 mutations have been implicated in cognitive and seizure disorders in young children^19,20^. JNK family kinases are well known for their role in the cellular stress response, and in neurons, they are capable of phosphorylating both scaffold and receptor molecules^21,22^ and serve as established regulators of neuronal differentiation and plasticity (for reviews see^23-25^). Here we have investigated the functional links between the disease-associated proteins SAP102 and JNK3, and we demonstrate that the behaviour of SAP102, in particular its mobility into and out of dendritic spines, is influenced by JNK signalling. We next demonstrated that kainate receptor trafficking can likewise by influenced by JNK regulation, and we showed that SAP102 forms a specific complex with JNK3 and the kainate receptor subunit GluK2; we thus provide insights into how these three proteins participate in a common molecular cascade that is disturbed in multiple monogenic neurodevelopmental disorders.

## Results

### SAP102 mobility is modulated by JNK activity

In immunofluorescence experiments, we observe colocalisation of endogenous JNK3 with the postsynaptic marker protein Homer in dendritic spines of mature primary neurons (Fig. 1A), confirming the presence of endogenous JNK3 at postsynaptic sites. We have previously shown that JNK3 binds to the postsynaptic scaffold protein SAP102^19^, and we observe overlapping expression of these two proteins in dendritic spines (Fig. 1B). In order to explore the consequences of the JNK-SAP102 interaction at synaptic sites, we generated a GFP-tagged SAP102 recombinant protein for analysis in living neurons.

**Figure 1:**
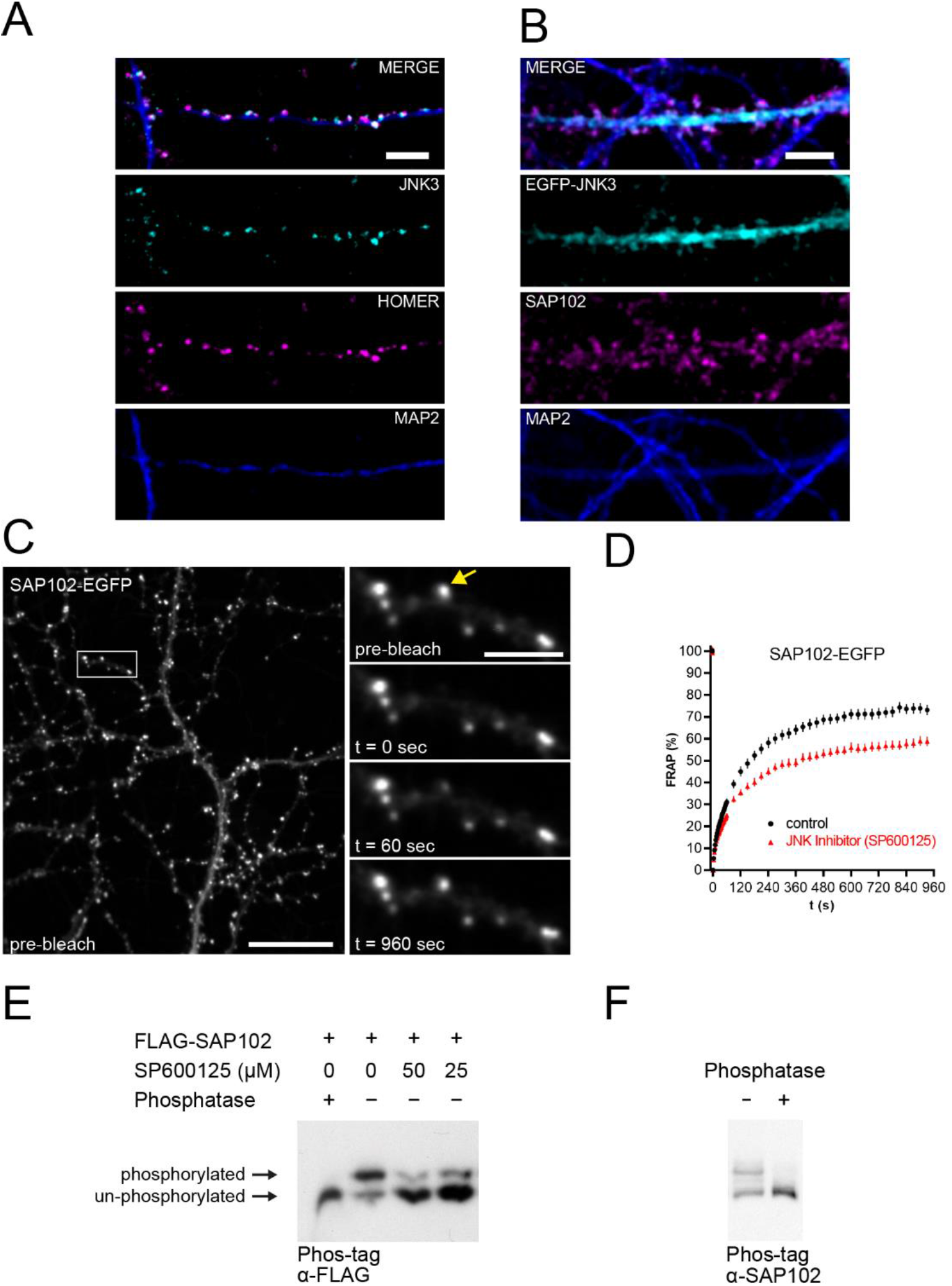
SAP102 mobility is modulated by JNK activity. **A**. Co-immunofluorescence of JNK3 (cyan, Alexa488) with synaptic marker protein Homer (magenta, Alexa568) in MAP2-positive dendrites (blue, Alexa405) of primary rat hippocampal neurons (DIV22), analysed by confocal microscopy. Scale bar: 5 µm. **B**. Co-immunofluorescence of lentiviral transduced and expressed EGFP-JNK3 (cyan, Alexa488) with SAP102 (magenta, Alexa568) in MAP2-positive (blue, Alexa405) dendrites of primary rat hippocampal neurons (DIV22), analysed by confocal microscopy. Scale bar: 5 µm. **C**. Live-cell imaging of SAP102-EGFP, following lentivirus-mediated gene delivery in primary rat hippocampal neurons (DIV23), using spinning disc microscopy for FRAP experiments: representative images of the same ROI with SAP102-EGFP in spines before photo-bleaching (pre-bleach), and at t=0 s, t=60 s, and t=960 s after photo-bleaching. Scale bar: 20 µm (overview) or 5 µm (zoom). **D**. FRAP experiments of SAP102-EGFP in spines showing control (black circle) vs. JNK inhibitor SP600125-treated (red triangle) rat primary hippocampal neurons (25 µM SP600125, 2 hours). Data are background subtracted, normalised and expressed as mean +/- SEM, with n=29-37 analysed spines from 3 images per condition. **E**. FLAG-SAP102 is phosphorylated in CHL cells. Phosphorylated proteins are separated from unphosphorylated proteins using Phos-tag-SDS-PAGE gels and analysed by western blot (α-FLAG). Phosphatase treatment shows unphosphorylated FLAG-SAP102 as control. Treatment of cells with JNK inhibitor (SP600125: 50 µM / 25 µM) results in a decrease of phosphorylated SAP102 protein. **F**. Endogenous SAP102 is phosphorylated in primary rat hippocampal neurons (DIV24); phosphatase treatment of cell lysate serves as control in Phos-tag/SDS-PAGE experiments followed by western blot (detection with α-SAP102).

SAP102 is a highly mobile scaffold protein, capable of moving rapidly into and out of dendritic spines^16^. Using fluorescence recovery after photobleaching (FRAP) of SAP102-EGFP in spines of rat hippocampal neurons (Fig. 1C), we were able to assess the effects of JNK modulation on SAP102 mobility. Following application of the JNK inhibitor SP600125, the mobile fraction of SAP102 is substantially reduced (Fig. 1D), indicating a functional link between SAP102 and JNK activity.

We next explored the idea that this functional link might reflect a direct JNK-mediated phosphorylation of SAP102. Using the Phos-tag system^26^, we analysed the phosphorylation status of SAP102: in CHL cells, we observed a distinct single phosphorylation of the overexpressed FLAG-tagged SAP102 (Fig. 1E upper band). The phosphorylated SAP102 protein (upper band) was completely absent after treatment with protein phosphatase, and the ratio of phosphorylated to unphosphorylated SAP102 was reduced in a dose-dependent fashion in response to treatment with a JNK inhibitor, suggesting that the observed SAP102 phosphorylation is indeed JNK-dependent. In line with this result, Phos-tag analysis of endogenous SAP102 in neuronal lysates indicates that a significant portion of the endogenous SAP102 protein is phosphorylated in mature cultured primary neurons (Fig. 1F). Together, these results indicate that SAP102 mobility is regulated by JNK activity, and they provide evidence in support of a direct JNK-mediated phosphorylation of SAP102 that may influence its behaviour.

### SAP102 is phosphorylated at position S368 by JNK

Next, we explored the putative phosphorylation of SAP102 by JNK in more detail. We analysed the SAP102 amino acid sequence for putative MAPK/JNK phosphorylation motifs and focussed on the proline-directed serine at position S368 of SAP102 that lies in close proximity to the predicted JNK docking site^27^ in a disordered region of the linker between the second PDZ domain and the MAGUK module PDZ3-SH3-GK of SAP102 (see scheme in Fig. 2A). Upon mutation of this serine to alanine, phosphorylation of the tagged SAP102 (as assessed by Phos-tag assay following expression in CHL cells) was no longer observed (see Supplementary Fig. S1), indicating that indeed the observed phosphorylation of SAP102 reflects phosphorylation at this site. To study this phosphorylation in detail, we raised phospho-specific antibodies against a peptide in which the serine at position 368 was phosphorylated. As expected, the affinity-purified antibody (α-p-SAP102) recognised the phosphorylated form of SAP102: in lysates from CHL cells overexpressing FLAG-tagged wild-type SAP102, we observed a single band at the expected size that was not present in lysates that had been treated with phosphatase (Fig. 2B). Importantly, the α-p-SAP102 did not recognise the overexpressed human SAP102 with S368A mutation (Fig. 2B), further confirming its specificity. This antibody was subsequently used to assess the phosphorylation of SAP102 at serine 368 by JNK3: in *in vitro* non-radioactive kinase assays, we observed a clear kinase-dependent accumulation of phosphorylated SAP102 (Fig. 2C).

**Figure 2:**
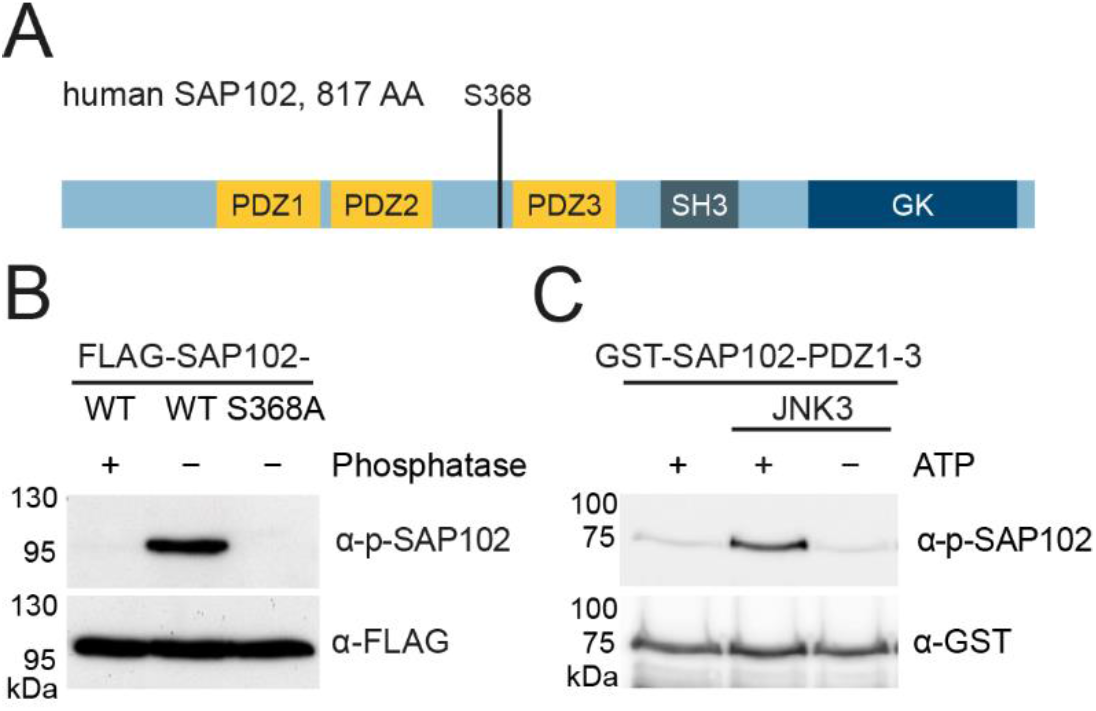
SAP102 is phosphorylated at S368 by JNK. **A**. Schematic overview of the human SAP102 MAGUK protein (UNIPROT #Q92796, 817 AA) with its annotated domains (PDZ1/2/3, SH3, and GK, drawn to scale) and phosphorylation site S368. **B**. FLAG-SAP102-WT and FLAG-SAP102 with phospho-deficient mutation S368A (FLAG-SAP102-S368A) were transfected in CHL cells and analysed by western blot for phosphorylation using the α-p-SAP102 antibody (raised against short peptide harbouring a phosphorylated S368). The dephosphorylated FLAG-SAP102 (phosphatase treatment) and the phospho-deficient mutation (FLAG-SAP102-S368A) are not detected by α-p-SAP102. α-FLAG detection serves as expression control of FLAG-SAP102 constructs. **C**. *In vitro* kinase assay of bacterially expressed and purified GST-SAP102-PDZ1-3 protein with active kinase JNK3 followed by western blot. α-p-SAP102 detects phosphorylated SAP102, α-GST serves as loading control for the GST substrate.

### JNK-mediated SAP102 phosphorylation is activity-dependent and developmentally regulated

We next investigated JNK-dependent phosphorylation of SAP102 in primary hippocampal neurons. Interestingly, in lysates from mature rat hippocampal neurons (DIV24), we observed a basal phosphorylation of SAP102 (Fig. 3A), but we did not detect this basal phosphorylation in immature neurons, i.e. prior to synaptogenesis (see Fig. 3C). This led us to hypothesise that JNK-mediated phosphorylation of SAP102 might be important in the regulation of synaptic connections, rather than in early development prior to synapse maturation.

**Figure 3:**
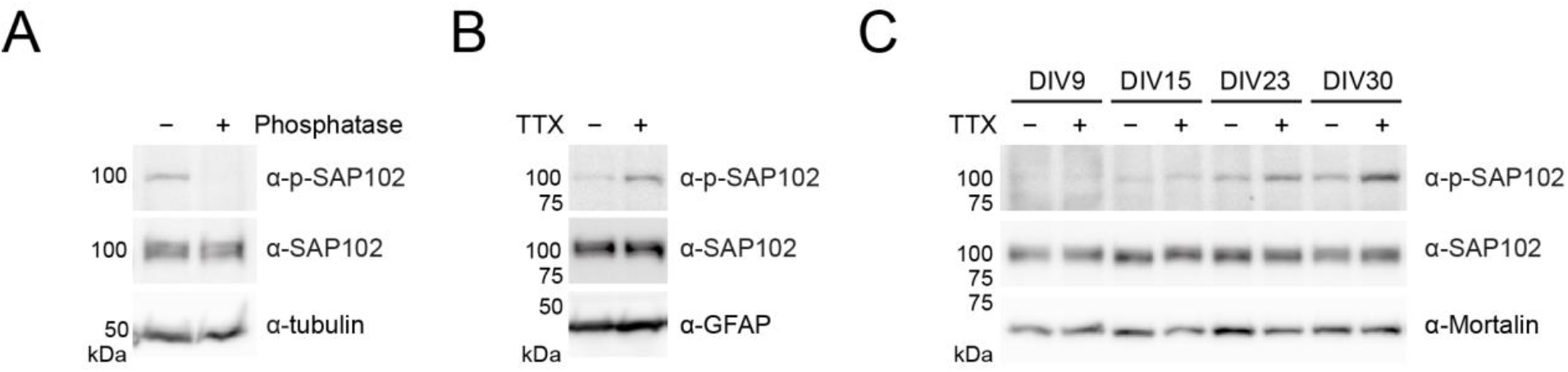
JNK-mediated SAP102 phosphorylation is activity-dependent and developmentally regulated. **A**. SAP102 is phosphorylated in primary rat hippocampal neurons. Lysates (DIV24) were analysed by western blot with the antibodies indicated; α-tubulin served as loading control. Phosphatase treatment of the lysate served as control for specificity of phosphoprotein detection. **B**. TTX treatment (2 µM, O/N) of primary rat hippocampal neuron cultures (DIV23) increases SAP102 phosphorylation, as observable by western blot with the phospho-specific SAP102 antibody (α-p-SAP102). α-SAP102 and α-GFAP served as controls. **C**. SAP102 phosphorylation in primary rat hippocampal neurons at DIV9, DIV15, DIV23, DIV30 was analysed by western blot following TTX treatment (2 µM, O/N). TTX-induced phosphorylation of SAP102 increases at later developmental stages (DIV23 and DIV30). α-SAP102 and α-Mortalin served as controls.

To explore the role of JNK-mediated phosphorylation of SAP102 in such dynamic processes, we induced synaptic upscaling (homeostatic synaptic plasticity) by activity blockade (blocking of Na^2+^ channels with TTX). Indeed, the phosphorylation of SAP102 at serine 368 strongly increased in mature neurons in response to TTX treatment (Fig. 3B, Supplementary Fig. S2i), suggesting that JNK might play a role in synaptic activity-dependent regulation of SAP102. In a subsequent set of experiments, we observed that this TTX-mediated phosphorylation of SAP102 was likewise developmentally regulated: When rat hippocampal neurons were analysed at different developmental time points for SAP102 phosphorylation (Fig. 3C), we detected basal phosphorylation of SAP102 at S368 after two, three and four weeks in culture (DIV15, DIV23 and DIV30), but not at DIV9. The phosphorylation of SAP102 at S368 increased strongly at DIV23 and even more at DIV30, when neurons were treated with TTX, whereas there was no clear TTX-induced increase of phosphorylation at DIV9 or DIV15. The observed increase in basal phosphorylation during development (compare lanes 1, 3, 5, and 7 in Fig. 3C) corresponds to the period of synapse formation, indicating a putative role for SAP102 phosphorylation at synapses. The idea that JNK-mediated regulation of SAP102 is part of the molecular response to changes in synaptic activity is further supported by the fact that we consistently observe a strong increase in phosphorylation during the process of synaptic upscaling, i.e. after TTX treatment in mature neurons.

Importantly, we observed this increase of SAP102 phosphorylation following TTX treatment not only in whole cell lysates from cultured rat hippocampal neurons but also in crude synaptosome fractions (Supplementary Fig. S2ii), suggesting that SAP102 is indeed phosphorylated when present in PSDs, which is in line with a role for JNK-mediated regulation of SAP102 at synaptic sites.

### Neuronal activity blockade and JNK inhibition exert opposing effects on SAP102 mobility

We have clearly shown that SAP102 is phosphorylated by JNK and that this phosphorylation is developmentally regulated and modulated by neuronal activity (Fig. 3). Given that JNK inhibition results in a decrease of both SAP102 phosphorylation (Fig. 2) and SAP102 mobility (Fig. 1), combined with the fact that SAP102 phosphorylation by JNK is activity-dependent, we hypothesised that SAP102 mobility might likewise be influenced by modulating neuronal activity.

We induced homeostatic plasticity with TTX-mediated activity blockade, and we subsequently analysed the mobility of SAP102 using FRAP (Fig. 4A). The mobile fraction of SAP102-EGFP indeed increased after induction of synaptic scaling, whereas it decreased following JNK inhibition. Simultaneous combination of the two treatments resulted in a compensation, yielding a mobility that was comparable to that for SAP102 in the untreated control condition (Fig. 4Ai). Together, these data indicated that JNK inhibition and TTX-mediated activity blockade have contrasting effects on SAP102 behaviour in this context, and highlight a putative role for JNK in coordinating the dynamics of the SAP102 function in dendritic spines, perhaps in concert with TTX-induced signal cascades that modulate homeostatic plasticity.

**Figure 4:**
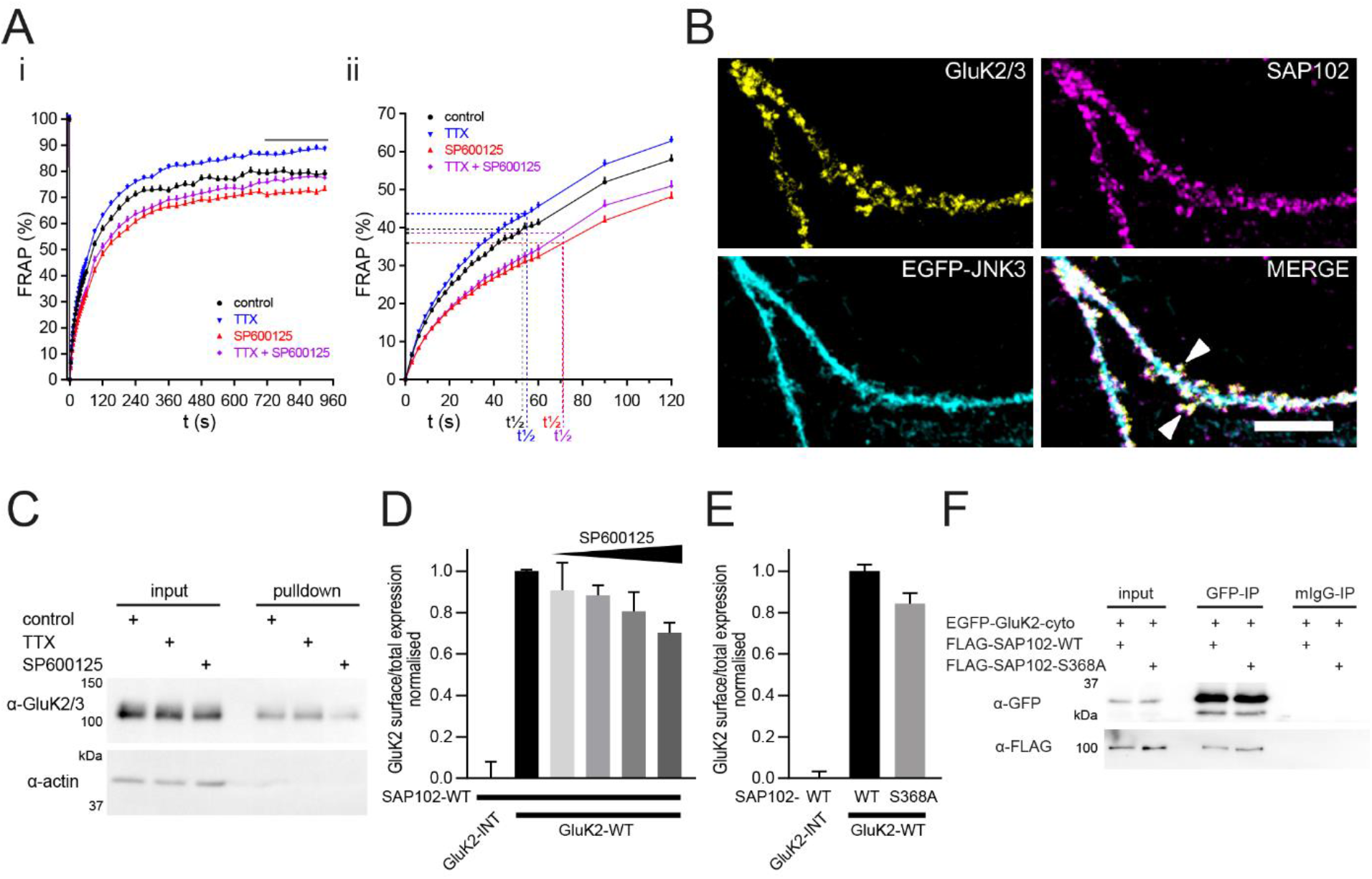
Both activity blockade and JNK activity promote mobility of SAP102 and GluK2 surface expression. **A**. FRAP experiments of SAP102-EGFP localised in spines of rat hippocampal neurons (DIV20-DIV23) were performed using live-cell imaging with spinning disc microscopy. Samples were treated with either TTX (2 µM, O/N), JNK inhibitor SP600125 (25 µM, 2 hours) or TTX and SP600125 (2 µM TTX O/N and 25 µM SP600125 2 hours prior to measurement). Data are background-subtracted, normalised (to the mean of seven pre-bleach acquisitions (=1) as well as to the value at t=0 s (=0)) and show the mean +/- SEM of n=94-112 spines (ROI) per condition (from 10-15 spines per image; 2-4 images per fluorodish; three fluorodishes from three different neuronal cultures). FRAP data (Ai) were acquired for 960 s. The mobile fraction was calculated as the mean of the last 8 FRAP values of SAP102-EGFP in each condition. Mobile fraction was used for estimation of half-time of recovery (t1/2) of mobile SAP102-EGFP (Aii), based on the FRAP experiments shown in Ai. **B**. Co-staining of endogenous GluK2/3 (yellow, Alexa405), SAP102 (magenta, Alexa568) and EGFP-JNK3 (cyan, EGFP/Alexa488) in rat hippocampal neurons (DIV22). Arrowheads show partial co-localisation of all three proteins in spines (MERGE). Scale bar: 10 µm. **C**. Surface biotinylation experiment in rat hippocampal neurons (DIV21) after O/N treatment with either TTX (2 µM) or JNK inhibitor SP600125 (25 µM) compared to control (DMSO). Proteins located at the cell surface were biotinylated and after pulldown with streptavidin-Dynabeads, surface expression of GluK2/3 was analysed by western blot (comparison of pulldown to input). The cytoplasmic protein actin (α- actin) serves as negative control for the surface biotinylation procedure (as expected, cytosolic proteins are not labelled). **D**. Analysis of GluK2 surface expression (relative to total GluK2 expression) following expression in CHL cells (On-Cell Western, OCW) together with wild-type SAP102 indicates that the relative GluK2 surface expression decreases with increasing amounts of JNK inhibitor SP600125. On the far left, the GluK2-INT variant serves as a control and reflects the zero value for GluK2 surface expression in this assay (see Supplementary Fig. S3 for comparison with cytosolic control protein). The black bar reflects the maximum GluK2 surface expression in this assay (cells are treated with a DMSO control; values are normalised to 1), and subsequent bars reflect comparable conditions after treatments with increasing inhibitor concentrations (25, 50, 100, and 200 µM). Data are mean +/- SD of a representative experiment (performed in triplicate), normalised to MYC-GluK2-WT. **E**. Analysis of GluK2 surface expression (relative to total GluK2 expression) using the same surface expression assay (OCW) enables comparison of wild-type (WT) and phospho-deficient SAP102 (S368A) with regard to their ability to promote GluK2 surface expression. Again, the GluK2-INT variant serves as a control to approximate the zero value for surface expression. Data are mean +/- SD of a representative experiment (triplicates), normalised to MYC-GluK2-WT. **F**. FLAG-tagged SAP102 (either wild-type or S368A phospho-deficient mutant as indicated) were coexpressed with the EGFP-tagged GluK2 cytosolic region in CHL cells. Following pulldown of the GluK2 C-terminus with the GFP antibody, coimmunoprecipitated SAP102 variants were analysed by western blot (α- FLAG), which indicated comparable interaction of the two variants with the GluK2 C-terminus. Mouse IgG served as negative control for the pulldown (mIgG-IP).

Importantly, not only the mobile fraction of SAP102-EGFP in spines, but also the t_1/2_ – which indicates the speed of recovery (half-recovery time) – was affected by JNK modulation. While TTX-treated neurons have a t_1/2_ of 55 seconds on average (which is very close to the t_1/2_ of 52 seconds observed in untreated neurons), JNK inhibition resulted in a significantly increased half-time of recovery (t_1/2_ of 71 seconds). Interestingly, this JNK-mediated reduction in the recovery speed was not altered by the additional application of TTX (combined treatment with TTX and SP600125 resulted in a t_1/2_ of 71 seconds), suggesting that JNK activity may act downstream in homeostatic mechanisms that mediate SAP102 accumulation at synapses.

In summary, these FRAP results show that JNK inhibition slows the speed of SAP102 movement into and out of dendritic spines. Moreover, in contrast to treatments that block activity and result in synaptic upscaling, JNK inhibition reduces the fraction of SAP102 molecules that are mobile. We conclude that JNK activity has a positive regulatory effect on SAP102 mobility, at least in part by influencing the speed at which SAP102 molecules are able to enter and/or exit the spine.

### SAP102 promotes KAR surface expression through formation of a functional tripartite complex with JNK

SAP102 has been shown to bind directly to glutamate receptors (*e.g*. NMDARs, KARs, and AMPAR auxiliary proteins). We therefore asked whether SAP102 – with the assistance of JNK – might be responsible for the shuttling of such receptors into or out of the spines during synaptic scaling. GluK2 is particularly interesting in the context of JNK regulation because there are similarities between the GluK2 and JNK3 knockout mice; moreover, patients with GluK2 or JNK3 mutations have overlapping phenotypic features^19,20,28^. Immunofluorescence analysis of the three proteins in primary hippocampal neurons fixed at DIV 21 indicated that they indeed exhibit overlapping localisation in dendritic spines (Fig. 4B). We next assessed the surface expression of endogenous GluK2 in surface biotinylation experiments (Fig. 4C). Following JNK inhibition in primary rat hippocampal neurons, a strong decrease in GluK2 surface expression was observed, whereas receptor surface expression after synaptic upscaling (TTX treatment) increased. These data are in line with a role for JNK-mediated regulation of the shuttling of kainate receptors to the surface. Together with our previous results demonstrating the JNK-mediated regulation of SAP102 mobility, this result suggests that SAP102 and JNK may work together to coordinate kainate receptor trafficking.

To investigate this hypothesis, we took advantage of an in-cell/on-cell western blot assay in heterologous cells, which enabled us to specifically examine the cooperative effects of JNK and SAP102 on kainate receptor subunit surface expression. Following expression of full-length GluK2 together with wild-type SAP102, we observed that a substantial fraction of overexpressed GluK2 was integrated into the membrane and thus detectable by staining prior to cell permeabilisation. For comparison, we took advantage of targeted GluK2 mutants that have been shown to remain internalised^29^. For these variants, expression ratios (surface:total) were comparable to ratios for overexpressed SAP102, which is entirely cytoplasmic (see Supplementary Fig. S3); we thus took advantage of this GluK2 variant to reflect our baseline for receptor surface expression in subsequent experiments.

We next assessed the effects of JNK inhibition on the surface expression of GluK2 in this assay. Following co-expression of SAP102 and GluK2 in CHL cells, treatment with JNK inhibitor SP600125 resulted in a decrease in the surface expression of the wild-type GluK2 in a dose-dependent fashion (Fig. 4E).

We then compared the effects of WT SAP102 with that of SAP102 harbouring a point mutation at the JNK phospho-site that precluded positive regulation by JNK (see SAP102 S368A depicted in Fig. 2A and corresponding Supplementary Fig. S1) on GluK2 surface expression in this assay. Compared to the wild-type, the SAP102 S368A mutant exhibited a reduced ability to promote the surface expression of the GluK2 subunit, suggesting that JNK-mediated phosphorylation of SAP102 indeed facilitates membrane expression of GluK2, as we hypothesised. In control experiments, we confirmed that both wild-type and phospho-deficient SAP102 mutants were comparably efficient in binding to GluK2 (see Fig. 4F), thereby excluding the possibility that differences in surface expression reflect reduced binding of the GluK2 C-terminus to SAP102 PDZ domains.

To further investigate this functional protein complex comprised of GluK2, SAP102, and JNK, we took advantage of coimmunoprecipitation assays with diverse combinations of recombinant proteins (see constructs depicted in Fig. 5A). We focussed initially on analysis of protein complexes that included JNK3 and GluK2. After expression of the full-length GluK2 together with JNK3, and pull-down of GluK2, we observed a clear coimmunoprecipitation of JNK3 (Fig. 5Bi). In subsequent experiments, we demonstrated that the GluK2 C-terminal cytoplasmic region (see schematic in Fig. 5A) is sufficient to enable binding to JNK3 (see Fig. 5Bii). As expected, this cytosolic region, which harbours a PDZ-binding motif at its distal C-terminus, also interacts with SAP102 via classical ligand-PDZ domain interactions that rely on the most C-terminal residues (see Supplementary Fig. S4; see also ^30-32^).

**Figure 5:**
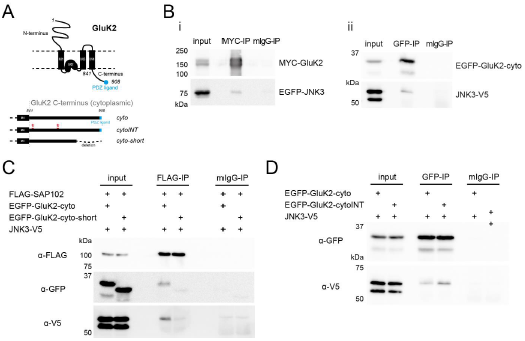
JNK3-SAP102-GluK2 protein complex. **A**. Schematic overview of the full-length kainate receptor subunit GluK2 (Uniprot P39087), depicting the extracellular N-terminus, the membrane-integrated domains (M1-4) and the intracellular, cytoplasmic C-terminus including the PDZ ligand (blue). Below an enlargement of the cytoplasmic GluK2 C-terminus is shown (drawn to scale), summarising the protein variants used in this study, including GluK2-cyto (with the wild-type PDZ-binding C-terminus), GluK2-cytoINT, which harbours mutations at two sites that are important for GluK2 receptor internalisation (highlighted in red), and GluK2-cyto-short, which reflects the wild-type C-terminus with a 26 AA deletion of the C-terminus. **B-D**. Interactions between of JNK3, GluK2 and SAP102 protein variants were analysed in co-immunoprecipitation experiments following expression in HEKT293 cells: proteins were precipitated using the antibodies indicated above, (mIgG served as negative control), and samples were analysed by western blot using antibodies as indicated. **B. i)** Pulldown of overexpressed full length MYC-GluK2 (MYC-IP) showed the co-immunoprecipitation with EGFP-JNK3. **ii)** Pulldown of the overexpressed cytoplasmic GluK2 C-terminus (EGFP-GluK2-cyto, GFP-IP) showed co-immunoprecipitation with JNK3-V5. **C**. Analysis of comparative co-immunoprecipitation experiments of overexpressed FLAG-SAP102 with JNK3-V5 and either EGFP-GluK2-cyto or EGFP-GluK2-cyto-short (deletion of PDZ ligand) indicated an increased tripartite complex formation when the cytoplasmic GluK2-C-terminus was intact (EGFP-Gluk2-cyto). **D**. Pulldown of either the wild-type GluK2 C-sterminus (EGFP-GluK2-cyto) or the internalised variant (EGFP-GluK2-cytoINT) with a GFP antibody (GFP-IP) enabled comparison of their ability to bind JNK3 (observable in western blot of coprecipated JNK3 with the α-V5 antibodies).

Interestingly, the presence of the wild-type GluK2 cytoplasmic region strengthens the interaction between SAP102 and the regulatory protein JNK3: when the full-length GluK2 C-terminus is expressed together with SAP102 and JNK3, the binding of JNK3 to SAP102 is clear, whereas when we delete the PDZ-binding motif of GluK2, the interaction between JNK3 and SAP102 is comparably weak (see Fig. 5C). We propose that binding of GluK2 to SAP102, which may be enhanced under specific physiological conditions, facilitates positive regulation by JNK and subsequent shuttling of GluK2 to the surface, with the assistance of the highly mobile SAP102. Analysis of targeted GluK2 mutants – namely, phospho-mimicking GluK2 point mutants that have been shown to remain internalised^29^ – provided insights into the physiological conditions that might favour such regulation by JNK. Such recombinant GluK2 C-termini (see schematic in Fig. 5A) display an increased binding affinity for JNK3 (Fig. 5D), suggesting that internalised GluK2 molecules might be especially available for JNK-mediated regulation, as one would expect if there was an acute physiological requirement for KARs to integrate into the synaptic membrane.

Our physiological observations in primary hippocampal neurons, together with our biochemical and pharmacological analysis in heterologous cells, strongly support the idea that JNK3, SAP102 and GluK2 are proteins that cooperate in the neuronal environment, and that formation of this multi-protein complex occurs in a controlled manner in response to specific signals, thus participating in the fine-tuned regulation of kainate receptor trafficking to the membrane.

## Discussion

In this study, we built on our previous observation that the postsynaptic scaffold protein SAP102 is biochemically associated with the regulatory kinase JNK3. Compared to related scaffold molecules, SAP102 is highly mobile, suggesting that it may play a unique role in trafficking receptors into and out of dendritic spines in response to physiological cues. We show here that this mobility is negatively regulated by JNK inhibition, and that SAP102 is indeed a direct phosphorylation target of JNK3. We further demonstrate that SAP102 and JNK3 are able to form a tripartite complex together with the ionotropic glutamate receptor GluK2. In line with a cooperative role for SAP102 and JNK3 in the regulated trafficking of this receptor, inhibiting JNK activity results in reduced GluK2 surface expression in cultured hippocampal neurons, which provides support for a model in which JNK-mediated regulation of SAP102 influences the trafficking of kainate receptors to postsynaptic sites (see model depicted in Fig. 6). Moreover, we demonstrate that the phospho-deficient SAP102 mutant, which cannot be phosphorylated by JNK, is less able than the wild-type to promote the surface expression of GluK2, which provides further evidence supporting a role for JNK-mediated regulation of SAP102 in coordinating GluK2 trafficking to the membrane.

**Figure 6:**
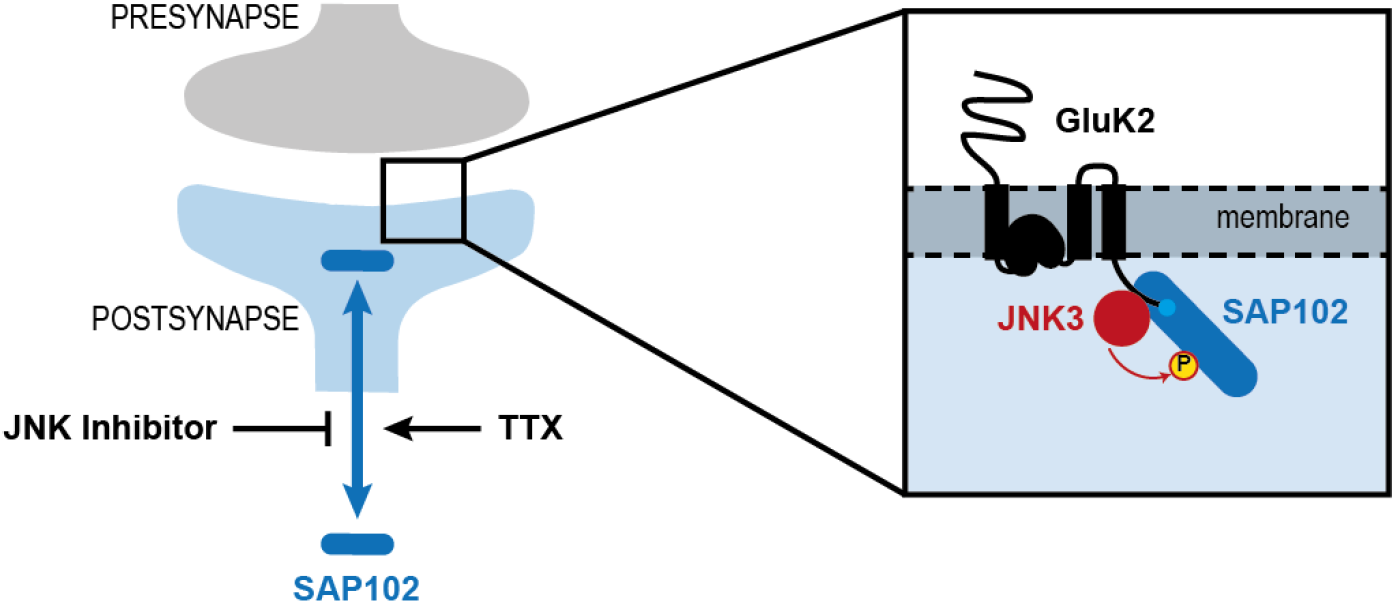
Model illustrating the functional interplay between JNK, kainate receptors and SAP102: JNK modulates kainate receptor surface expression via phosphorylation-dependent regulation of SAP102 mobility.

SAP102 mutations have been implicated in intellectual disability^11-15^ (see also OMIM #300850), and mice that lack the protein entirely exhibit defective spatial learning and aberrant synaptic plasticity^33^. It has been shown that SAP102, like PSD-95 and other synaptic MAGUKS, is of critical importance for regulating glutamate receptor trafficking to synaptic sites (*e.g*. ^34^), and the consequences of MAGUK loss are especially apparent during synaptogenesis and synapse maturation^35^, presumably in response to defective trafficking and anchoring of glutamate receptors and other PDZ binding transmembrane proteins at the PSD. Protein composition of the PSD of glutamatergic synapses is clearly important for normal network development^36,37^, and MAGUK proteins are central hubs within this scaffold; factors that affect their functional integrity and proper regulation are thus obvious targets for investigation if we want to understand the mechanistic basis of neurodevelopmental disorders. Interestingly, it has been shown that diverse post-translational modifications are indeed able to influence MAGUK function and resulting protein composition of the PSD^6,18,22,38^.

In this context, our observation that JNK activity modulates SAP102 mobility is of special relevance. JNK proteins are historically known for regulation of the cellular stress response and – in neurons – for modulating excitotoxicity. More recent studies have highlighted their importance in the regulation of normal physiological processes during neural development (for review see ^23^). In modulating the motion of SAP102 into and out of dendritic spines, JNK signalling can fine-tune the cellular localisation of this scaffold and its binding partners, which inevitably influences the protein composition of PSDs at glutamatergic synapses. Importantly, our observation that JNK inhibition reduces kainate receptor surface expression reveals one of the many downstream effects that could result from such hindered SAP102 mobility.

Also relevant in the context of our observations is the fact that the kainate receptor subunit GluK2, like JNK3 and SAP102, has been implicated in monogenic forms of cognitive and developmental disorders^28,39^. The GluK2 KO mouse exhibits deficits in learning^40^ and aberrant social and cognitive behaviour^41^, underscoring the importance of normal kainate receptor function for network formation and cognitive development. In our experimental setup, reduced JNK activity corresponds to reduced KAR expression at the membrane; the observed JNK-mediated effects on GluK2 provide a direct mechanistic link between the monogenic cognitive disorders associated with GluK2 and JNK3 mutations, and it is plausible that disease pathophysiology in both disorders involves related dysregulation of kainate receptor function. This functional link likewise provides a putative explanation for the common phenotype of the GluK2 and JNK3 KO mouse models: both are resistant to kainate-induced seizures^42,43^. Our data, in which we highlight functional and biochemical links between both of these proteins and the synaptic scaffold SAP102, support the idea that these three molecules participate in a common signal cascade coordinating the cellular response during kainate-induced excitotoxicity and in the regulation of kainate-receptor function under normal physiological conditions. Indeed the three proteins are able to form a complex, and our biochemical experiments suggest that the simultaneous presence the three proteins positively influences their individual affinities for one another: specifically, we observe an increased interaction stability between SAP102 and JNK3 in the presence of the PDZ-binding ligand GluK2 (see Fig. 5C). This result, which supports the idea that these three proteins indeed cooperate functionally, is in line with studies on macromolecular complex formation dictated by other MAGUKs: complex formation induced by ligand binding to PSD-95 PDZ domains has been demonstrated in both *in vitro* and cellular contexts^44-47^.

Finally, our data provide support for the idea that JNK3 participates in the *regulated* control of kainate receptor cycling to and from the postsynaptic membrane. We highlight that JNK inhibition leads to reduced KAR surface expression, and that normal JNK activity levels correspond to higher KAR surface expression. We also observe that JNK3 binds favourably to GluK2 variants mimicking those that are internalised in response to PKC-mediated phosphorylation^29,48,49^), in line with a regulatory action of JNK3 that is influenced by the cellular environment and the state of GluK2. We propose a functional model that reflects these data (see Fig. 6): in response to elevated synaptic activity, kainate receptors are post-translationally modified by other kinases, including *e.g*. PKC, and receptor internalisation is facilitated^50,51^, which prevents KAR hyperactivation and excitotoxicity. JNK3 binds preferentially to such internalised variants (Fig. 5D), perhaps in order to restore surface expression of synaptic KARs and maintain activity in a homeostatic process. Such a model depicts a positive role for JNK in modulating neuronal activity and likewise illuminates a putative mechanism for the documented resistance to kainate-induced seizures observed in both the JNK3 and GluK2 KO mice. Our data also support the idea that JNK3 executes its regulatory control of KAR function in cooperation with SAP102, which becomes more mobile when JNK is active or under conditions of reduced synaptic activity (see Fig. 1 and Fig. 4). Our model thus highlights how these three proteins participate in a common signalling pathway, highlighting a putative mechanistic explanation for some of the common behavioural and synaptic transmission aberrations found in the corresponding disease models.

## Methods

### Antibodies

Antibodies used in this study include FLAG (mouse, Sigma F1804), FLAG-HRP (Sigma A8592, WB 1:1000-4000), GFP (mouse, Roche, 11814460001; goat, Abcam, AB6673, WB 1:4000; chicken, Abcam, ab13970, IF 1:5000-7500, WB 1:5000), GST (goat, GE Healthcare, 27457701V, WB 1:5000), MYC (mouse, Clontech, 631206; rabbit, Cell Signaling, 2272S, WB 1:5000), V5 (rabbit, Millipore, AB3792, WB 1:5000) and those directed against the following proteins: Actin (rabbit, Sigma, A2066, WB 1 : 2000), GFAP (mouse, Antibodies Inc., 75-240, WB 1:2000), GluK2/3 (rabbit, Millipore, 04-921, WB 1:4000, IF 1:500), Homer (guinea pig, Synaptic Systems, 160004, IF 1:500), JNK3 (rabbit, Millipore, 04-893, IF: 1:250, WB 1:3000), MAP2 (mouse, IF 1:500; guinea pig, Synaptic Systems, 188004, IF 1 : 1000), Mortalin (mouse, Antibodies Inc., 75-127, WB 1:5000), SAP102 (mouse, Antibodies Inc, 75-058, WB 1:1000-5000, IF 1:100), SAP102 (rabbit, Abcam, ab3438, IF 1:500), Tubulin (rat, Abcam, ab6160, WB 1:12000).

For coimmunoprecipitation experiments, 2 μg of the respective antibody was used. Unspecific mouse IgGs (mIgG), as required (SantaCruz, SC-2025, 2 μg), were used for negative controls in coimmunoprecipitation studies. All primary and secondary antibodies were diluted in 5 % milk/PBST for western blot (WB) or in 4 % BSA/PBS for immunofluorescence (IF) experiments. For WB experiments, we used the following secondary antibodies: α-mouse-HRP (Dianova #115035003 or α- native-mouse-IgG-HRP Abcam, ab131368), α-rabbit-HRP (Dianova #111-035-003), or α-goat-HRP (Santa Cruz #sc-2020) with a dilution of 1:5000. α-rat-HRP (Santa Cruz #sc-2032) and α-chicken-HRP (Abcam #ab6753) were used with a dilution of 1:10000. For IF experiments, we used secondary antibodies α-guinea-pig-Alexa405 (Abcam, ab175678), α-chicken-Alexa488 (Dianova #703-545-155) and the following antibodies from Thermo Fisher Scientific: α-mouse-Alexa405 (A-31553), α-rabbit-Alexa405 (A-31556), α-rabbit-Alexa488 (A-21441), α-guineapig-Alexa568 (A-11075), α-mouse-Alexa568 (A-11031), α-rabbit-Alexa568 (A-11036), all with a dilution of 1:1000. For On-Cell Western experiments, we used α-mouse-680RD (1:500, Li-COR #926-68070) and α-mouse-800CW (1:500, Li-COR #925-32218).

### Antibody against phosphorylated-S368-SAP102

The phosphorylation site S368 of SAP102 lies in the linker region between PDZ2 and PDZ3 of human SAP102 (DLG3_HUMAN, Q92796, 817 AA; OMIM #300850); S368 of human SAP102 corresponds to position S386 in rat SAP102 (DLG3_RAT, Q62936, 849 AA). Customised polyclonal phospho-specific antibodies were raised in rabbit against a phospho-peptide including pS368 (DLG3_HUMAN, AA 361-373: QVPPTRY-S*(PO3H2)*-PIPRH; EUROGENTEC) and affinity-purified (α-p-SAP102, rabbit, WB: 1: 1000-1500).

### DNA constructs

The construct pEGFP-C1-JNK3a2 has been published previously^19^ and was used as a template for cloning JNK3 into pBudCE4 under the control of the EF1a promotor to generate a C-terminal V5 tagged expression construct JNK3-V5 (pBudCE4-JNK3-V5). We used the pEGFP-C1-JNK3 as a template for cloning EGFP-JNK3 into the lentiviral shuttle vector f(syn)w (based on FUGW^52^) for expression under the control of a synapsin promotor in primary rat hippocampal neurons.

Phospho-deficient SAP102 variant (pCMV2A-FLAG-SAP102-S368A) was generated by site-directed mutagenesis using pCMV2A-FLAG-SAP102^19^ as a template. Full-length SAP102 was also cloned into pCMV3A to generate N-terminal MYC-tagged construct (pCMV3A-MYC-SAP102). For the generation of the SAP102 lentiviral expression construct, we cloned SAP102 into the pEGFP-N1 vector using pCMV2A-FLAG-SAP102 as a template. In a second step, we used the pEGFP-N1-SAP102 as a template for cloning SAP102-EGFP into the lentiviral shuttle vector f(syn)w. The N-terminal GST-SAP102-PDZ1-PDZ3 fusion construct (pGEX-6P-1-FLAG-SAP102-PDZ1-3) was subsequently generated from pCMV2A-FLAG-SAP102. This fragment, encoding an N-terminal FLAG tag and the first 472 amino acids of SAP102, was cloned into the pGEX-6P-1 vector with an N-terminal GST tag (GE Healthcare).

The construct pcDNA3-6xMYC-GluK2 was a gift from C. Mulle (Bordeaux, France) and was used as a template for the cloning of the cytoplasmic C-terminal tail of GluK2 (pEGFP-C1-GluK2-cyto; 841-908 aa of P39087, GRIK2_MOUSE), the shorter deletion construct of the cytoplasmic tail of GluK2 (pEGFP-C1-GluK2-cyto-short; 841-882 aa of P39087) and the generation of mutant variants by site-directed phospho-mimicking mutagenesis of S846E and S868E (pEGFP-C1-GluK2-cytoINT, pcDNA3-6xMYC-GluK2-INT).

### Cell culture

CHL-V79 cells and HEK293T cells were maintained in DMEM supplemented with 10 % FBS, 2 mM L-glutamine and penicillin/streptomycin in a humidified incubator at 37 °C with 5 % CO_2_. Transfections were performed using Lipofectamine 2000 (Invitrogen) and Opti-MEM (Gibco) according to manufacturers instructions.

Primary rat E18 hippocampal neurons were prepared as already described^53^. Briefly, embryonic E18 Wistar rats were used for hippocampi isolation and dissociation. Dissociated cultures were plated in Neurobasal medium, infected at DIV10 with lentivirus and harvested, treated or analysed after three weeks in culture between DIV20 and DIV24. Treatments of neurons with JNK inhibitor SP600125 (25 µM) or TTX (2 µM) were done in conditioned medium.

### Immunofluorescence and confocal microscopy

Immunofluorescence experiments were performed as described previously^19^ according to standard IF protocols. Dissociated cultures of primary rat hippocampal neurons were fixed in 4 % PFA in PBS for 10 min, washed in PBS, permeabilised in 0.2 % Triton-X in PBS for 5 min, washed in PBS, blocked in 4 % BSA in PBS for 1 hour at room temperature. Cells were incubated with primary antibodies over night at 4 °C in blocking solution, washed with PBS, incubated with secondary antibodies for 30-60 min in blocking solution and finally washed in PBS. Coverslips were mounted on glass slides with Fluoromount-G (Southern Biotech).

Cells were imaged using a confocal laser scanning microscope (TCS-SP5 II, Leica). Images were acquired with a 63X objective (1.5-2x zoom, 1024×1024 px, 0.4 µm steps in z, z-planes). Images shown are single z-planes with the strongest signal intensity in the region of interest.

### Live cell imaging: Fluorescence Recovery After Photobleaching (FRAP)

Primary rat E18 hippocampal neurons plated in Fluorodishes (cover-glass bottom, 35 mm, WPI Inc.) were infected at DIV10 and analysed in live cell experiments at DIV20-24. Neurobasal medium was exchanged with Tyrode solution (25 mM HEPES pH 7.4, 120 mM NaCl, 2.5 mM KCl, 2 mM CaCl_2_, 2 mM MgCl_2_, 30 mM Glucose) 30 min before measurement. In case of treatment with SP600125 or TTX, conditioned medium including the drugs was exchanged with Tyrode solution including the respective treatment.

FRAP experiments were performed using a NIKON spinning disc confocal CSU-X microscope (FRAP/PA, 60x Plan Apo objective with NA = 1.4, laser: 488 nm, emission filter: 525/50, Andor DU-888 X-9798 camera, Perfect Autofocus PFS, NIS Software). Fluorodishes were acclimated in the live cell chamber of the microscope for 20 min (37 °C, 5 % CO_2_) before starting the measurement. Spines (10-15 spines per image) of SAP102-EGFP expressing neurons were selected as ROIs for bleaching and imaging: 10 pre-bleach images, photo-bleaching (Laser 488 nm 100%, dwell time 100 µs, 15x bleaches), followed by 20 acquisitions with an interval of 3 s and further followed by 30 acquisitions with an interval of 30 s (post-bleach images: Laser 488 nm 20%, 300 ms exposure time, 538×538 px, total duration of FRAP measurement 960 sec). Three FRAP acquisitions per dish have been performed sequentially.

Images have been analysed using NIKON NIS Software for ROI inspection over time, drift correction in X/Y, fluorescence intensity measurements and background subtraction. Spines, which disappeared during measurement, were excluded from further analysis. ROIs of highly moving spines were manually adjusted. For the FRAP calculation, the first three of ten pre-bleach images were deleted ^54^ and the fluorescence values for each ROI over time were background-corrected and normalised (EXCEL): the mean of the last seven of ten pre-bleach values was set to 1 (100 %), whereas the lowest intensity value (first post-bleach image, t=0) was set to 0 (0 %). All other intensity values were normalised accordingly for each ROI. We measured per condition (control, TTX, SP600125, TTX+SP600125, Fig. 4) a total of 94-112 spines (10-15 spines per image; 2-4 images per fluorodish; three fluorodishes from three different neuronal cultures) and calculated the mean and the SEM (GraphPad Prism 7). The mobile fraction was calculated as the mean of the last eight FRAP values for each condition. The t_1/2_ (half-time of recovery) as the half maximal recovery time of the mobile fraction of SAP102-EGFP was estimated on the basis of the FRAP graph.

### Co-immunoprecipitation

Transfected cells (CHL V79 or HEKT293) were harvested 20 to 45 hours after transfection and resuspended in lysis buffer (50 mM Tris-HCl pH 7.5, 100 mM NaCl, 0.1-1% Triton-X, including protease inhibitors (Mini Complete without EDTA, Roche)). After lysis using a 30-gauge syringe and 30 min incubation on ice, lysates were centrifuged twice (10 min, 20000 *g*, 4 °C) and the resulting supernatant was incubated with 2 µg antibody or normal mouse IgG for 3 hours at 4 °C in a spinning wheel. To remove precipitated proteins, the lysates were centrifuged (10 min, 20000 *g*, 4 °C) and the resulting supernatants were added to 25 µl Protein G Agarose (Roche), rotating for 1 hour at 4 °C, followed by three times washing (each 5-10 min) with lysis buffer. Analysis of precipitated were analysed by western blotting.

### In vitro kinase assay

The non-radioactive, *in vitro* kinase assay was performed as described before^19^. Briefly, bacterially expressed GST-SAP102-PDZ1-3 was purified, eluted, desalted ^53^ and subsequently used as a substrate for the phosphorylation by JNK3 (commercially available from BioMol). Reactions were performed in 20 mM Tris pH 7.5, 10 mM MgCl_2_, 200 μM ATP, 0.1 % beta-mercaptoethanol for 30 min at 30 °C. SAP102 phosphorylation was analysed by western blotting with α-p-SAP102.

### Surface Biotinylation

Primary rat hippocampal neurons (DIV20-24, in 6w) were washed with PBS on ice and then biotinylated for 30 min (1 mg/ml EZ-Link Sulfo-NHS-LC-Biotin, ThermoScientific, in PBS) on ice. After three PBS washing steps, the biotinylation reaction was quenched with 100 mM Glycine/PBS (2 x 5 min) and washed again twice in PBS on ice. Cells were harvested by scraping in lysis buffer (50 mM Tris-HCl pH 7.5, 100 mM NaCl, 1 % Triton-X, 0.1 % SDS, 0.5 % sodium deoxycholate including Mini Complete Inhibitors (Roche)) and lysed on ice for 30 min after sonication. Cleared lysate (twice centrifugation for 10 min @ 20000 g) was incubated for three hours with 50 µl Dynabeads M-280 Streptavidin (Invitrogen) followed by three washing steps with lysis buffer. Biotinylated proteins were analysed by western blotting.

### Western blot, Phos-tag

Protein samples for western blotting were boiled at 95 °C for 5 min in 2X SDS sample buffer and separated on standard Lämmli SDS-PAGE gels. Proteins were semi-dry blotted onto PVDF membranes (Roche), which were subsequently blocked in 5 % milk in PBST, incubated with primary antibodies in 5 % milk in PBST over night at 4 °C, followed by three PBST washes, incubation with the secondary antibodies for 1 hour in 5 % milk in PBST and final three washes with PBST. Chemiluminescence signals were detected using the imager ImageQuant LAS4000mini (GE Healthcare).

Phos-tag (WAKO) gel electrophoresis for analysis of phosphorylated SAP102 as well as the sample preparation for phosphatase treatment was done as described before^53^ using Bis-Tris-buffered neutral-pH gels with 6 % polyacrylamide supplemented with 20-75 µM Phos-tag.

### On-Cell Western

GluK2 surface expression experiments were performed using the cell-based On-Cell Western assay, that enabled the quantitative detection of surface expressed MYC-tagged GluK2 in heterologous CHL-V79 cells by an immunofluorescence staining procedure (near-infrared fluorescence) followed by acquisition using the LI-COR Odyssey CLx Imager. Transfected CHL cells in 48w plates were carefully washed in PBS, fixed in 4 % PFA-PBS for 10 min, washed in PBS, blocked for 30 min in BSA-PBS, incubated with α-MYC (mouse, 1:500, Clontech #631206) in 4 % BSA-PBS for 60 min at room temperature to stain surface expressed GluK2, washed in PBS, incubated with α-mouse-800CW (1:1000, LI-COR #925-32218) for 30 min and washed in PBS (surface staining completed), followed by a standard immunofluorescence protocol: samples were fixed in 4 % PFA-PBS for 10 min, washed in PBS, permeabilised for 5 min in 0.2 % TritonX-PBS, washed in PBS, blocked in 4 % BSA-PBS for 30 min, incubated with α-MYC (mouse, 1:1000) over night at 4°C, washed in PBS, incubated with α-mouse-680RD (1:1000, LI-COR #926-68070) for 30 min followed by PBS wash.

Experiments were performed in sextuplicates (six transfected 48wells per condition): three of the six wells were used for surface (800 nm) and total (700 nm) staining of MYC-GluK2 (triplicates), whereas the other three wells were used for controls of FLAG-SAP102 expression (800 nm) and cell density (700 nm). The staining procedure for the SAP102 expression and cell density control was performed as described above, but the surface staining steps were omitted. FLAG-SAP102 proteins were stained using α-FLAG (mouse, 1:1000) and α-mouse-800CW (1:1000), cell density was controlled by using CellTag700 (LI-COR, #926-41090) during the incubation of the secondary antibodies according to manufacturer’s recommendations.

The 48w plates were imaged using the LI-COR Odyssey CLx Imager and further analysed by using the software Image Studio (LI-COR Biosciences), FIJI^55^ (including the plugin Readplate2.1^56^), EXCEL (Microsoft) and Prism 7 (GraphPad). A grid was used to identify the wells, the mean grey values were measured for images at 700 and 800 nm, followed by a background subtraction for each channel. The ratio of the 800nm/700nm intensities (Surface GluK2 fraction/total GluK2) were calculated for each well. As a control for non-surface GluK2 expression we used the expression of an internalised GluK2 receptor variant (MYC-GluK2-INT with phospho-mimicking mutation at S846E and S868E). The 800/700 ratio (mean of triplicates) of the MYC-GluK2-INT samples was subtracted (set to 0) from all single wells followed by a normalisation to the mean of the triplicates of MYC-GluK2 surface(800)/total(700) staining (set to 1). Two to three experiments were performed in triplicate; data from representative experiments are shown.

## Acknowledgements

We are grateful for technical assistance from Melanie Fuchs, and for the support of the Virus Core Facility (VCF), the Advanced Medical Bioimaging Core Facility (AMBIO) and the Microscopy Core Facility (MCF) at the Charité-Universitätsmedizin Berlin. We are appreciative of ongoing input and structural support from both Dietmar Schmitz and Craig Garner. Finally, we are grateful for diverse project funding from the DFG (individual projects 261102178 and 431572356, as well as collaborative research grants EXC 257, CRC 665, CRC 958, EXC 2049). Both Hanna L. Zieger and Bettina Schmerl were additionally supported financially through the PhD fellowship program of the Charité-Universitätsmedizin Berlin.

