## Supplements for "JNK activity modulates postsynaptic scaffold protein SAP102 and kainate receptor dynamics in dendritic spines"

### Supplementary Figure S1

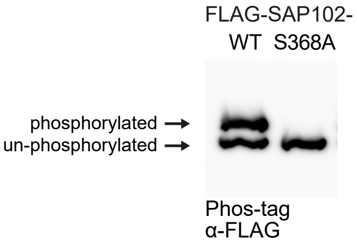


FLAG-SAP102 is phosphorylated at position S368 in CHL cells. Phosphorylated proteins are separated from unphosphorylated proteins using Phos-tag-SDS-PA-GE gels and analysed by western blot. Phospho-deficient point mutation at position S368A of FLAG-SAP102 showed only one band.

### Supplementary Figure S2

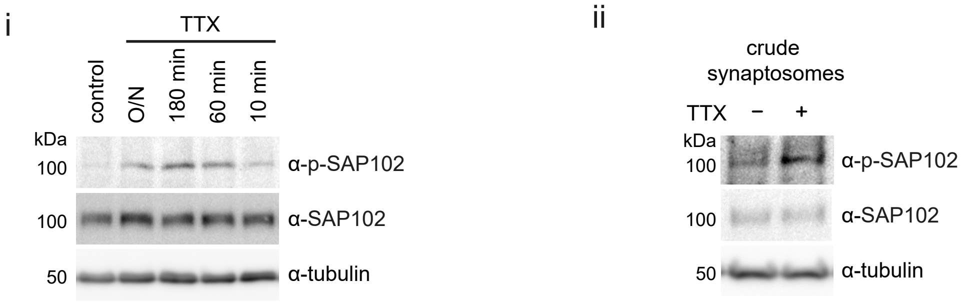


Endogenous neuronal SAP102 is phosphorylated after TTX treatment.
i) Western blot analysis of SAP102 phosphorylation in primary rat hippocampal neurons with 2 μM TTX treatment for different durations (O/N, 180 min, 60 min, 10 min); α-tubulin serves as a loading control. ii) Detection of phosphorylated SAP102 in crude synaptosome preparations from primary rat hippocampal neurons (DIV23).

### Supplementary Figure S3

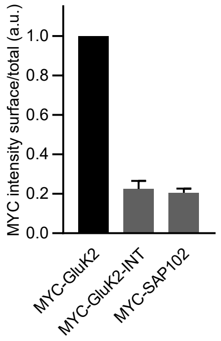


On-cell Western (OCW) assay for surface expression of overexpressed MYC-GluK2 in transfected CHL cells.
Normalised MYC surface fraction/total staining for MYC-GluK2-WT, MYC-GluK2-INT (internalised GluK2) and MYC-SAP102 (control: only cytoplasmic expression). Data are mean +/- SD of three independent experiments each consisting of technical replicates.

### Supplementary Figure S4

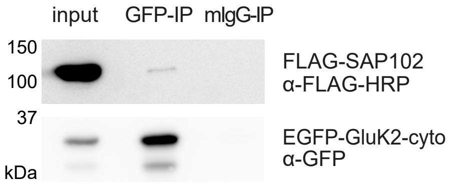


SAP102 interacts with the cytoplasmic GluK2-C-terminus in co-immunoprecipitation experiments (HEK293T). Pulldown of overexpressed EGFP-GluK2-cyto (GFP-IP) showed co-precipitation of FLAG-SAP102 compared to mIgG-IP (pulldown with unspecific mIgGs as negative control).
